# Experimenting with Two Recent Feature Selection Methods for High-Dimensional Biological Data

**DOI:** 10.1101/2025.09.09.675248

**Authors:** Minzhe Zhang, Xiao Yang

**Affiliations:** PromptBio Inc., 7068 Koll Center Pkwy, Suite 402, Pleasanton, CA 94566, USA

## Abstract

Feature selection in high-dimensional biological data, where the number of features far exceeds the number of samples, has long posed a significant methodological challenge. This study evaluates two recently developed feature selection methods, Stabl and Nullstrap, under a simulation framework designed to replicate regression, classification, and non-linear regression tasks across varying feature dimensions and noise levels. Our results demonstrate that Nullstrap consistently outperforms Stabl and other benchmarked methods across all evaluated scenarios. Furthermore, Nullstrap proved significantly faster and more scalable in high-dimensional settings, underscoring its suitability for large-scale omics data applications. These findings establish Nullstrap as a robust, accurate, and computationally efficient feature selection tool for modern omics data analysis.

## 1 Introduction

Feature selection is a critical step in the analysis of high-dimensional biological data, such as that generated by genomics, transcriptomics, and proteomics studies. Its primary goal is to identify a subset of relevant features associated with a specific outcome of interest, for instance, discovering biomarkers for disease status, drug response, or patient prognosis. However, the high-dimensional setting—where the number of features (*p*) far exceeds the number of samples (*n*)—poses a significant challenge. This “curse of dimensionality” often leads to model overfitting, reduced interpretability, and computational inefficiency, thereby complicating the reliable identification of robust biological signals.

Traditional feature selection methods can be broadly categorized into three groups: univariate filtering methods (e.g., mutual information, correlation, ANOVA, AUC), embedded methods (e.g., random forest, Lasso), and wrapper methods (e.g., forward selection, backward elimination) [1]. While these approaches have been widely used in the analysis of omics data, there is no single method that is universally accepted as the optimal choice for high-dimensional biological datasets, and the selection of an appropriate method often depends on the specific characteristics of the data and the analysis goals.

The development of the machine learning feature selection module for our AI-driven bioinformatics platform, PromptBio, has necessitated a critical evaluation of existing methods against these practical challenges. Consequently, we defined a set of criteria for an ideal feature selection method: (1) availability of mature and well-maintained implementations in common data analysis languages (such as Python and R) to ensure ease of use and reproducibility; (2) computational efficiency and scalability to handle large-scale datasets; (3) support for major supervised learning tasks relevant to biological research, including regression, classification, and survival analysis; (4) interpretability, specifically the ability to assign meaningful feature scores that reflect the influence of each feature on the outcome, with a preference for sparse solutions where most features receive a score of zero, akin to Lasso regularization [2]; (5) minimal reliance on manual hyperparameter tuning, achieved through internal mechanisms or heuristics that automate parameter optimization and reduce the demand for user expertise; and (6) an automatic and unbiased approach for determining or recommending the optimal number of features to select, rather than relying on subjective visual inspection or arbitrary thresholds.

Our search for a suitable method led us to two recently developed approaches: Stabl [3] and Nullstrap [4], which show significant promise in addressing these challenges. Both approaches are grounded in penalized linear modeling frameworks, such as Lasso and Elastic Net, and are specifically designed to address the challenge of false discovery rate (FDR) control through different null data generation strategies. Stabl controls the false discovery proportion (FDP) by augmenting the original dataset with artificially generated features, either via random permutation of the original features or through the use of Model-X (MX) knockoffs [5]. With the integration of noise injection and a data-driven signal-to-noise threshold, it enables more robust feature selection. Nullstrap generates synthetic null data by fitting a null model under the global null hypothesis—assuming that none of the features are associated with the outcome—without modifying the original data. In this study, we employ a simple simulation framework to compare their performance against baselines approaches. A summary comparing the key characteristics of both methods is provided in Supplementary Table S1. Our goal is to deliver a practical assessment and actionable guidance for applying Stabl and Nullstrap to large-scale biological data analysis.

## 2 Methods

### 2.1 Simulation Study

We generated synthetic data to evaluate the performance of feature selection methods. The simulation framework creates feature matrices with known correlation structures and target variables with controlled signal-to-noise ratios.

#### 2.1.1 Feature Matrix Generation

Let 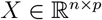 be a feature matrix generated from a multivariate normal distribution:

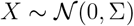

where *n* is the number of samples, *p* is the number of features, and Σ is a correlation matrix obtained from real biological data (Supplementary Information 1). This correlation matrix captures the inherent dependency structure present in high-dimensional biological datasets.

#### 2.1.2 Regression Target Variable Generation

For regression tasks, the target variable *y*_reg_ is generated using a linear model with additive noise:

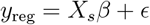

where 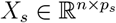 represents the subset of signal features (the first *p*_*s*_ features),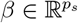 is a coefficient vector with entries sampled uniformly from [−1, −0.5] ∪ [0.5, 1], and *ϵ* ∼ 𝒩 (0, *σ*^2^) is Gaussian noise with standard deviation *σ* = *α* · std(*X*_*s*_*β*). The noise level parameter *α* controls the signal-to-noise ratio.

#### 2.1.3 Classification Target Variable Generation

For classification tasks, we first generate the continuous output *y*_cont_ using the same linear model as in regression, then transform it to binary labels. The continuous output is transformed using a logistic function applied to the centered values:

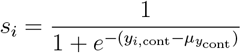

where 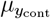 is the mean of *y*_cont_ across all samples, and *s*_*i*_ represents the transformed probability score for sample *i*.

The binary classification labels are determined by applying a threshold *τ* to these transformed scores. For each simulation run, *τ* is randomly sampled from a uniform distribution *τ* ∼ Uniform(0.3, 0.7), and the initial binary labels are computed as:

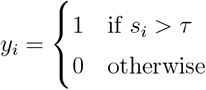

To simulate imperfect labels, we introduce controlled noise by randomly flipping a subset of the generated labels. Let 𝒩 ⊂ {1, 2, …, *n*} be a randomly selected subset of indices with cardinality |𝒩 | = ⌊*α* · *n*⌋, where *α* ∈ [0, 1] is the noise ratio parameter. The classification labels with noise are then computed as:

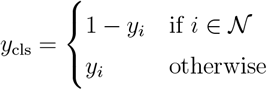

To ensure balanced classification datasets, we implement a post-generation quality check on the final noisy labels. If the resulting ratio of either class (0 or 1) falls outside the range [0.25, 0.75], the threshold *τ* is regenerated and the entire classification process (including noise introduction) is repeated until the class balance requirement is met.

#### 2.1.4 Non-linear Regression Target Variable Generation

For non-linear regression tasks, the target variable *y*_nl_ is generated using a feedforward neural network architecture with ReLU activation functions. The network consists of *L* hidden layers, where each layer transforms the input through a series of non-linear operations.

Let *h*^(0)^ = *X*_*s*_ be the initial input layer containing the signal features, and let 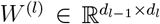 be the weight matrix for the *l*-th hidden layer, where *d*_*l*_ is the dimension of the *l*-th hidden layer. The forward propagation through the network is defined recursively as:

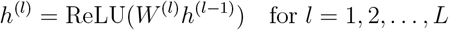

where the ReLU activation function is defined as ReLU(*x*) = max(0, *x*).

The weight matrices *W* ^(*l*)^ are generated using the same strategy as the coefficient vector *β* in the linear regression case, with entries sampled uniformly from [−1, −0.5] ∪ [0.5, 1]. The final output layer produces the continuous target variable:

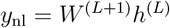

where 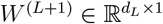 is the output layer weight matrix.

This approach allows us to generate target variables with non-linear dependencies on the signal features, where the degree of non-linearity can be tuned by adjusting the neural network architecture (e.g., number of layers or hidden units).

#### 2.1.5 Simulation Settings

The simulation framework supports various configurations with different sample sizes, feature dimensions, and noise levels. In our experiments, we focused on scenarios with *n* = 300 samples and *p*_*s*_ = 10 signal features containing information for target prediction. For regression tasks, we set the number of features to *p* ∈ {1000, 3000, 10000} and the noise ratio to *α* ∈ {0.2, 0.4, 0.6}. For classification tasks, we used *p* = 1000 features and a noise ratio of *α* ∈ {0.0, 0.05, 0.1}. For non-linear regression tasks, we considered two neural network architectures: one with a single hidden layer of *d*_1_ = 8 units, and another with two hidden layers, each with *d*_1_ = *d*_2_ = 8 units, to model varying complexity. For both classification and non-linear regression tasks, we fixed the number of features at *p* = 1000 to simplify the evaluation. The impact of feature dimensionality on performance was assessed in the regression experiments.

### 2.2 Benchmarking Methods

Both Stabl and Nullstrap methods were applied to classification and regression tasks—including both linear and non-linear settings—using their default parameters (Supplementary Information 2). These methods generated feature ranking scores and selected feature sets for subsequent evaluation.

For baseline comparison, we implemented two widely-used feature selection approaches: a mutual information-based filtering method and a random forest method. In the mutual information method, we calculated the mutual information between each feature and the response variable, resulting in a score for each feature. For the random forest method, we trained a random forest model and obtained feature importance scores from the fitted model (Supplementary Information 2). In both cases, these scores were used to rank the features for subsequent selection. To facilitate a direct and consistent comparison, we selected the top 10 features from mutual information and random forest methods to form the baseline feature sets.

### 2.3 Evaluation Metrics

We evaluate the performance of feature selection methods using two complementary metrics: Area Under the Precision-Recall Curve (AUPRC) and F1 score.

AUPRC measures the overall ranking quality of features, where informative features should be ranked higher than non-informative ones. Given the extreme class imbalance in our simulation (most features are non-informative), AUPRC is preferred over Area Under the Receiver Operating Characteristic curve (AUROC) as it focuses specifically on positive class performance and is not dominated by the majority negative class.

F1 score provides a balanced measure of precision and recall, evaluating the final feature set selected after applying a threshold. While AUPRC assesses ranking quality, F1 score measures the practical utility of the selected feature subset in real-world applications where a definitive feature set is required.

We also record wall-clock time for the core feature selection procedure to compare computational costs across methods. Given the variability in implementation details (e.g., hyperparameter search grids, bootstrap iterations, parallelization strategies), these comparisons provide general computational cost estimates for practical usage rather than comprehensive algorithmic efficiency analysis.

## 3 Results

Across all simulation scenarios, Nullstrap consistently outperformed Stabl, random forest, and mutual information-based feature selection methods. In regression tasks, Nullstrap achieved the highest AUPRC and F1 scores, particularly in settings with low noise and lower feature dimensionality, where it nearly perfectly identified informative features (Figure 1). As the number of non-informative features or the noise level increased, the performance of all methods declined, but Nullstrap maintained a clear advantage over the alternatives. Stabl generally ranked second, followed by random forest, while mutual information exhibited the weakest performance throughout. We also observed that as noise increased, Nullstrap’s F1 score declined as expected, while Stabl’s F1 score remained relatively stable. This is likely due to Stabl’s intrinsic procedure for optimizing the frequency threshold used in feature selection.

**Figure 1:**
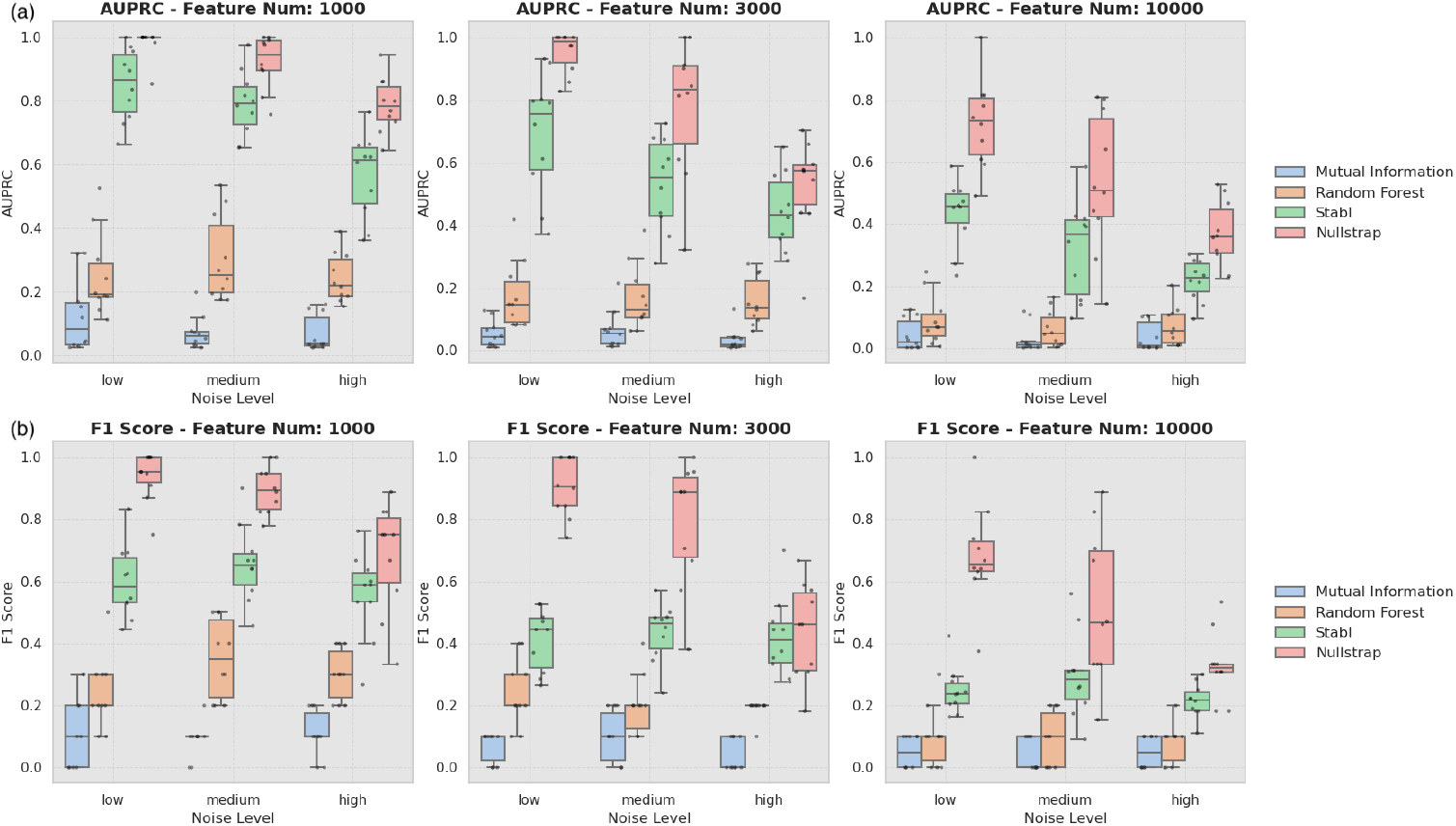
Performance of feature selection methods in simulated linear regression tasks. Performance is shown as (a) AUPRC and (b) F1 score across simulation settings with feature dimensions *p* = 1000, 3000, and 10 000, and noise levels corresponding to *α* = 0.2 (low), 0.4 (medium), and 0.6 (high).

Similar trends were observed in classification (Figure S2) and non-linear regression tasks (Figure S3): Nullstrap remained the top-performing method, with Stabl and random forest trailing behind and mutual information performing the worst. Overall, feature selection was most effective in linear regression, followed by classification, and was most challenging in non-linear regression. The performance gap between Nullstrap and Stabl narrowed as task complexity increased, but Nullstrap consistently provided superior feature ranking and selection across all evaluated conditions.

In terms of computational cost, Nullstrap was significantly faster than all other methods, including the univariate mutual information feature ranking approach. In our simulations, the time required by Nullstrap, mutual information, and random forest methods increased roughly in proportion to the number of features (see Supplementary Table S2). By contrast, Stabl was not only the most computationally expensive method, but its computation time increased more rapidly than the growth in feature dimensionality, making it less scalable to high-dimensional datasets.

## 4 Discussion

This simulation study systematically benchmarked two recently proposed feature selection methods, Nullstrap and Stabl, against established baselines in high-dimensional, biologically realistic scenarios. Our results demonstrate that both Nullstrap and Stabl outperform traditional approaches such as random forest and mutual information filtering, with Nullstrap consistently achieving the highest accuracy in both feature ranking and selection. Notably, Nullstrap maintained its superior performance even in challenging settings characterized by high noise levels, increased dimensionality, and non-linear relationships between features and outcomes.

A key strength of Nullstrap is its computational efficiency, enabling rapid analysis of large-scale omics datasets. In contrast, while Stabl is competitive in terms of feature selection accuracy and offers the advantage of adaptively choosing the frequency threshold to control the FDR, it exhibited substantially higher computational costs and poorer scalability as the number of features increased. This difference in computational burden is particularly relevant for modern biological studies which often involve datasets with tens of thousands of features or more.

It is important to acknowledge that, although our simulation framework incorporates realistic aspects of biological data, it does not capture the full complexity of real-world omics datasets. Both Nullstrap and Stabl rely on penalized linear models as their core feature selection engine, which aligns closely with the linear and sparse structure of our simulated data. Future work should explore the performance of these methods in settings with more complex non-linearities, or alternative data-generating mechanisms to further assess their generalizability.

In summary, our findings highlight Nullstrap as a robust, accurate, and scalable feature selection method for high-dimensional biological data. Its strong performance across a range of simulated scenarios, combined with its computational efficiency, makes it a practical and reliable tool for researchers seeking to identify informative features in modern omics analyses.

## Acknowledgements

We gratefully acknowledge Vincent Guo and Shishir Hebbar for their contributions to the literature review of feature selection methods.

## Supplementary Information

### Supplementary Information 1: Correlation Structures in Simulated Data

To create realistic correlation structures for simulated data, we used gene expression data from the CPTAC endometrial cancer dataset, specifically from Supplementary Table S2 of [6]. Genes with very low (*<*0.5) or very high (*>*5) variance were removed. We then calculated pairwise correlations among the remaining genes. To reduce redundancy, one gene from each highly correlated pair (absolute correlation *>* 0.95) was removed, keeping the gene with the lower index. The distribution of pairwise correlation coefficients in this dataset after filtering are shown in Figure S1. For each simulation, we randomly sampled the required number of features (e.g., *p* = 1000, 3000, 10 000) from this filtered set to generate the simulated datasets.

### Supplementary Information 2: Method Implementation Details

Implementation details for each feature selection method are provided below. For Stabl and Nullstrap, we used either default settings or the example code from the respective package documentation.

- **Mutual Information:** Feature scores were computed using the `mutual_info_regression` and `mutual_info_classif` functions from the `sklearn.feature_selection` Python module.
- **Random Forest:** Feature importance was computed using the `RandomForestClassifier` and `RandomForestRegressor` functions from the `sklearn.ensemble` Python module. The following hyperparameters were tuned using 5-fold cross-validation: `n_estimators` (20, 50, 100, 200), `max_depth` (None, 5, 10), and `min_samples_split` (2, 5, 10). The best parameters were selected and the model was retrained on the entire dataset before computing feature importances.
- **Stabl:** For regression, we used `Lasso(max_iter=int(1e6))` as the base estimator; for classification, we used `LogisticRegression(penalty=“l1”, max_iter=int(1e6), solver=“liblinear”)` . Stabl was run as `Stabl(base_estimator=clone(lasso), lambda_grid=“auto”, verbose=1)` with default frequency threshold optimization.
- **Nullstrap:** For classification, we used `fit <- nullstrap_filter(X, y, fdr value = 0.1, best_lambda = NULL, B reps = NULL, dist type = “normal”, model_type = “glm”)` ; for regression, `model_type = “linear”` was used.

## Supplementary Figures

**Figure S1:**
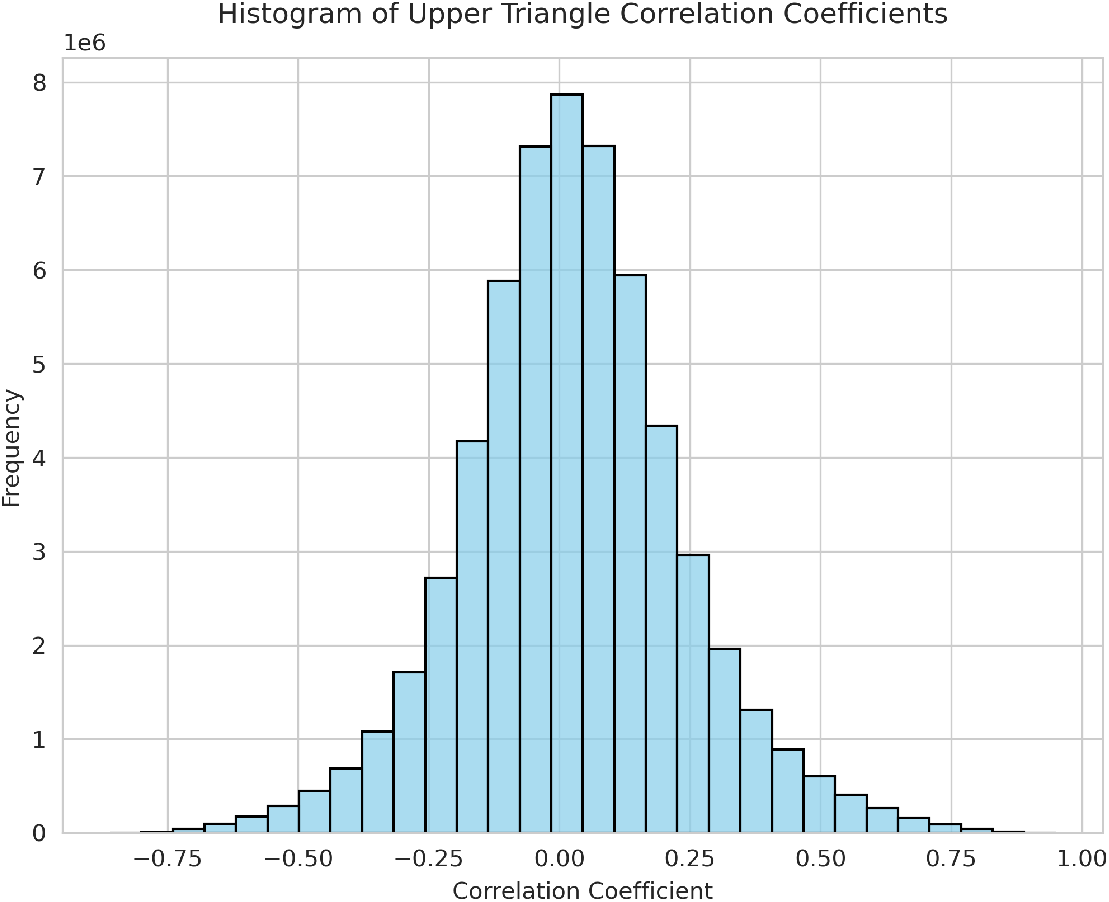
Distribution of pairwise correlation coefficients in the filtered CP-TAC endometrial cancer gene expression dataset. The histogram shows the density of correlation coefficients among all pairs of genes after variance and redundancy filtering.

**Figure S2:**
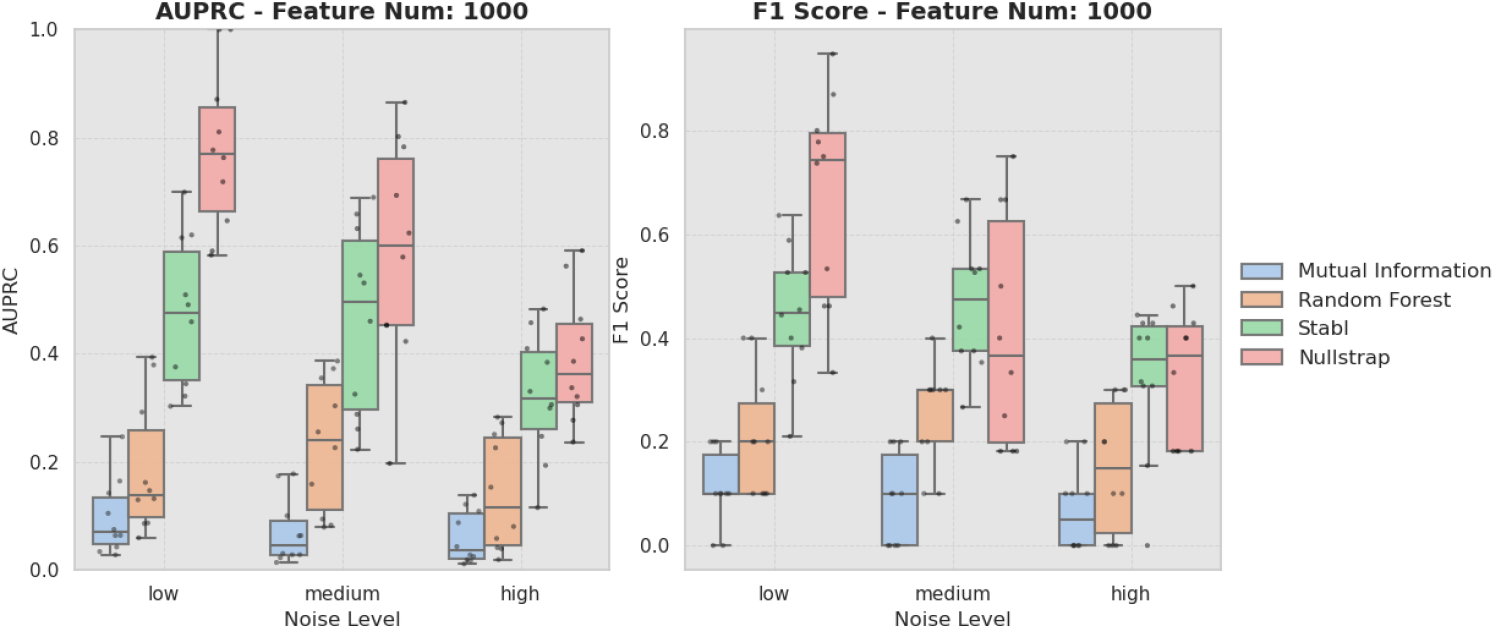
Performance of feature selection methods in simulated classification tasks. Performance is shown as AUPRC and F1 score for the classification task with feature dimension *p* = 1000 and noise levels corresponding to *α* = 0.0 (low), 0.05 (medium), and 0.1 (high).

**Figure S3:**
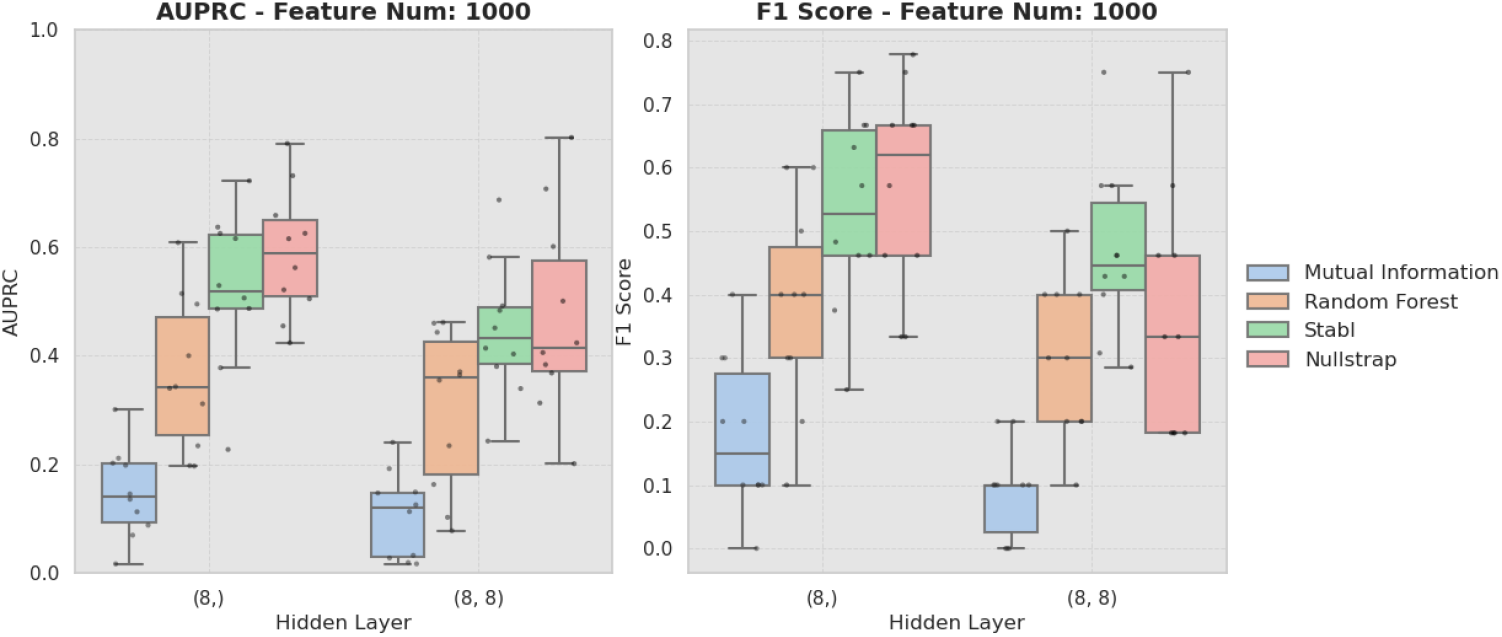
Performance of feature selection methods in simulated non-linear regression tasks. Performance is shown as AUPRC and F1 score for the non-linear regression task with feature dimension *p* = 1000 and neural network architectures with 1 hidden layer (8 units) and 2 hidden layers (8, 8 units).

## Supplementary Tables

**Table S1:**
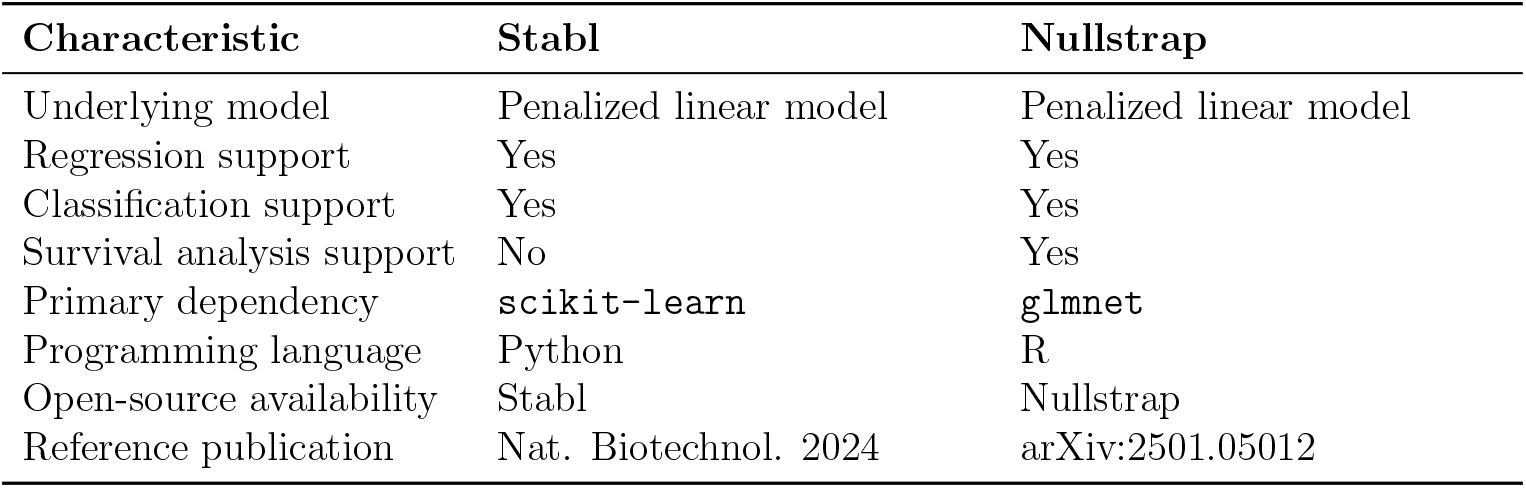
Comparison of Stabl and Nullstrap feature selection methods. Summary of key characteristics, supported analysis types, software dependencies, programming language, and availability for the Stabl and Nullstrap methods.

**Table S2:** Computation time (in seconds) for feature selection methods across simulation settings. Mean wall-clock time (± standard deviation) is reported for each method, averaged over 10 simulation replicates. Results are shown for linear regression tasks with varying feature dimensions (*p*).

| Method | $p = 1000$ | $p = 3000$ | $p = 10\,000$ |
| --- | --- | --- | --- |
| Mutual Information | $1.96 \pm 0.02$ | $5.87 \pm 0.05$ | $19.52 \pm 0.19$ |
| Random Forest | $41.03 \pm 0.77$ | $122.16 \pm 1.90$ | $406.27 \pm 7.78$ |
| Stabl | $92.60 \pm 2.13$ | $348.66 \pm 29.11$ | $1644.52 \pm 151.70$ |
| Nullstrap | $1.17 \pm 0.16$ | $3.08 \pm 0.40$ | $8.50 \pm 0.75$ |

